# HMOX1 controls a heme–ferritin switch that protects cells from ferroptosis

**DOI:** 10.1101/2025.10.14.682408

**Authors:** Izadora de Souza, Izabela Amélia Marques Andrade, Florencio Porto Freitas, Ana Beatriz da Silva Teixeira, Maria Carolina Clares Ramalho, Karoline Almeida Lima, Ancely Ferreira dos Santos, Werner Schmitz, Gholamreza Fazeli, Clarissa Ribeiro Reily Rocha, José Pedro Friedmann Angeli

## Abstract

Modulating the intracellular labile iron pool (LIP) has emerged as a promising strategy to induce ferroptosis in cancer cells, offering a way to overcome resistance to apoptosis-based therapies. One of the main contributors to LIP is heme catabolism mediated by heme oxygenase-1 (HMOX1), which promotes ferroptosis sensitivity by releasing free iron. Beyond its role as an iron donor, heme can influence diverse proteins and signaling pathways that drive tumor progression, but how heme regulates ferroptosis remains poorly understood. Here, we uncover a paradoxical, protective function of heme in the absence of HMOX1 activity. When HMOX1 is inactive, heme becomes stabilized, leading to ferritin upregulation, suppression of ferroptosis, and rescue of cell death induced by both pharmacological and genetic inhibition of GPX4. Our findings reveal an unrecognized heme–HMOX1–ferritin axis that controls ferroptosis sensitivity. Targeting this pathway may offer a new therapeutic strategy to modulate ferroptosis in cancer.

## Introduction

Ferroptosis is a distinct, iron-dependent form of cell death characterized by the uncontrolled accumulation of lipid hydroperoxides within cellular membranes. Unlike apoptosis or necrosis, ferroptosis is triggered when a lipid repair enzyme that detoxifies phospholipid hydroperoxides glutathione peroxidase 4 (GPX4) is inhibited or inactivated. GPX4 normally detoxifies phospholipid hydroperoxides by reducing them to non-toxic lipid alcohols. In its absence, lipid hydroperoxides accumulate, destabilize membranes, and trigger cell death.

The central driver of this process is the Fenton reaction, in which ferrous iron Fe(II) reacts with hydrogen peroxide (H_2_O_2_) to generate highly reactive oxygen radicals, including hydroxyl radicals (HO•). These radicals abstract a bis-allelic hydrogen atom from polyunsaturated fatty acids (PUFAs) within phospholipids, generating carbon-centered radicals. The latter react with molecular oxygen (O_2_) to form lipid peroxyl radicals (HOO•), which perpetuate chain reactions by extracting hydrogen atoms from nearby acyl chains^1^. Under physiological conditions, GPX4 stops this cascade, thereby maintaining membrane integrity. However, when GPX4 is inactivated, uncontrolled iron-driven lipid peroxidation disrupts the cell membrane, leading to cell death^2,3^.

Because cancer cells maintain higher intracellular iron levels than non-malignant cells, they are believed to be especially susceptible to ferroptosis ^4,5^. Cancer cells tightly regulate iron homeostasis to ensure that enough iron is available for essential cellular processes, including enzyme function, heme synthesis, and iron-sulfur cluster assembly. This balance also minimizes the risk of iron-mediated oxidative damage ^6-9^, and if it is disrupted, the intracellular labile iron pool (LIP) increases, sensitizing cells to ferroptosis ^10^. In this context, manipulating iron metabolism has emerged as a potential strategy to treat drug-resistant tumors. One major contributor to LIP is the degradation of free heme, an iron-containing protoporphyrin complex. This process is catalyzed by heme oxygenase-1 (HMOX1), releasing carbon monoxide, biliverdin, and ferrous iron. The release of iron through this pathway directly contributes to ferroptosis sensitivity ^11-14^.

However, a great deal of evidence has shown that the role of heme in cancer biology and ferroptosis extends beyond serving as an iron donor. Heme regulates transcription, translation, drug metabolism and signal transduction pathways that affect tumor growth and proliferation ^15-18^. Despite this, the biological relevance of heme in cancer and ferroptosis has often been attributed exclusively to its iron content, overlooking potential regulatory functions mediated by the intact heme molecule.

In the present study, we explored the regulatory role of heme in ferroptosis in cancer cells and uncovered an unexpected cytoprotective function. We found that in the absence of HMOX1, heme suppressed ferroptosis by upregulating ferritin expression. The effect required the loss of the catalytic activity of HMOX1, rather than its nuclear localization, and occurred independently of the BACH1/NRF2 pathway, a key regulator of antioxidant defense. In addition, we discovered that stabilization of heme preserved cell viability even in cells without GPX4, a condition that normally produces a lethal phenotype with increased susceptibility to ferroptosis. This indicates that loss of HMOX1 strongly suppresses ferroptosis. Mechanistically, we observed that stabilization of heme, combined with an iron-sequestration response caused by inactivation of iron regulatory protein 2 (IRP2), a key promoter of iron uptake under low-iron conditions, leads to increased expression of the ferritin heavy and light chains. Collectively, these findings uncover a previously unrecognized ferroptosis-suppressive role of heme in the absence of HMOX1, mediated in part by ferritin induction.

## Results

### 1. Hemin protects HMOX1-knockout cells from ferroptosis

By degrading heme and releasing ferrous iron, HMOX1 contributes to the labile iron pool (LIP) and thereby increases ferroptosis sensitivity (Fig. 1a). To further investigate the link between heme accumulation and ferroptosis sensitivity, we used CRISPR-Cas9 gene editing to knock out HMOX1 (HMOX1^KO^) in A375, A549, and HT1080 cells. We confirmed successful HMOX1 knockout by immunoblotting. HMOX1 expression in these cells was induced by adding hemin, the oxidized and more stable form of heme, which is widely used experimentally to mimic free heme exposure and induce HMOX1 expression. As expected, immunoblotting revealed strong HMOX1 expression in control cells (sgNT) but not in HMOX1^KO^ cells (Fig. 1b, Extended Data 1a). Uncontrolled heme exposure can cause cell damage as it exerts pro-oxidant effects ^19^ by catalyzing reactive oxygen species (ROS) formation and promoting lipid peroxidation. Consistent with this, we observed that HMOX1 deficiency strongly sensitized the cells to high doses of hemin (0 - 128 μM) (Fig. 1c, Extended Data 1b). A non-toxic concentration of hemin (10 μM), combined with GPX4 inhibitor ML210, a ferroptosis inducer, enhanced ferroptosis sensitivity in control cells. Strikingly, however, the same treatment rendered HMOX1^KO^ cells markedly more resistant to ferroptosis, producing the opposite effect (Fig. 1d, Extended Data 1c,d). This unexpected result reveals that the absence of HMOX1 in the presence of hemin paradoxically protects against ferroptosis.

**Fig. 1.**
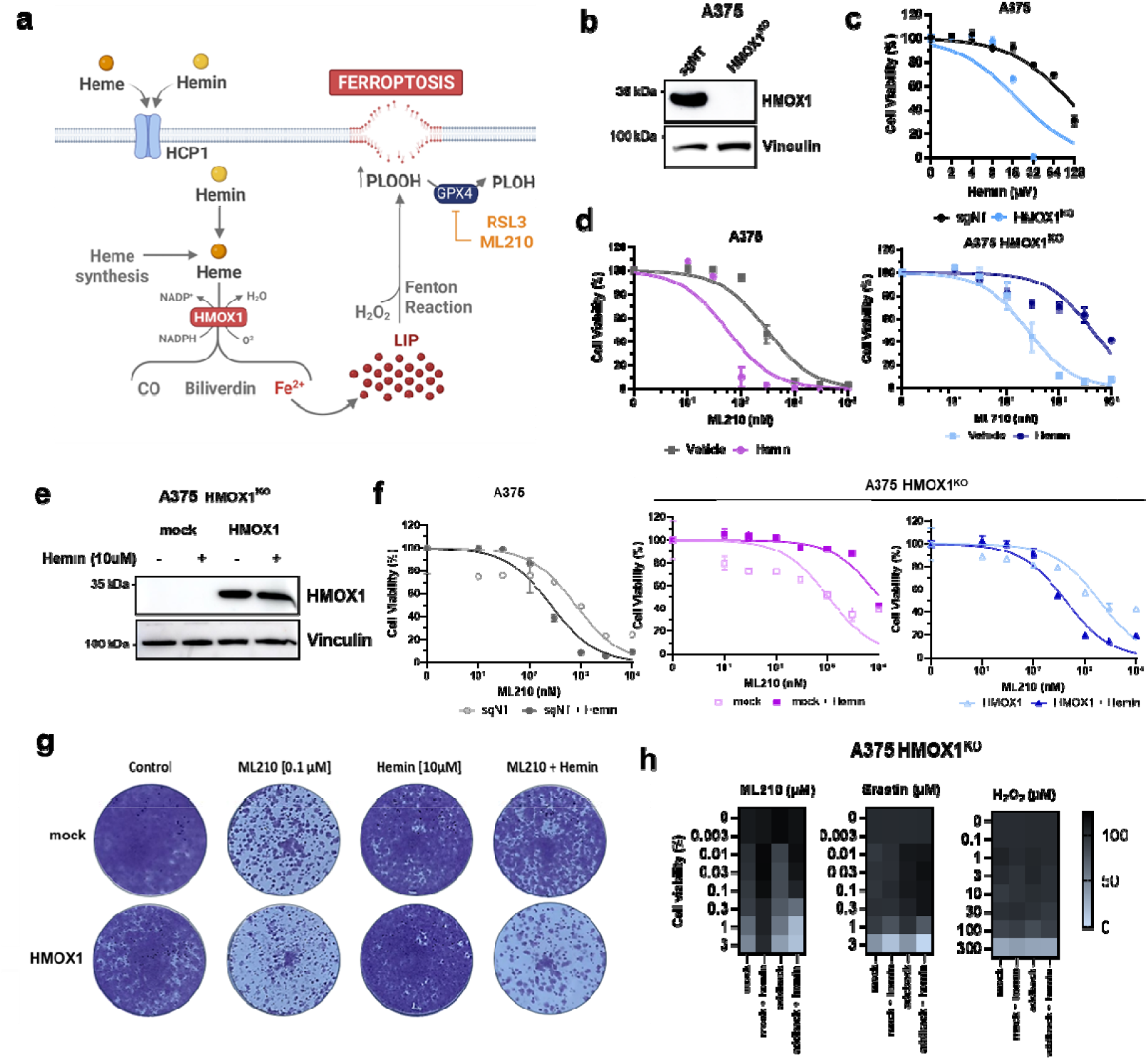
Hemin protects HMOX1-knockout cells from ferroptosis. **a**, Schematic view of the known role of hemin in ferroptosis. Heme synthesized in mitochondria through sequential enzymatic steps is exported to the cytosol, where it can be degraded by heme oxygenase-1 (HMOX1) into biliverdin, carbon monoxide (CO), and ferrous iron (Fe2□). The released iron contributes to the labile iron pool (LIP), promoting lipid peroxidation and ferroptotic cell death. Under ferroptosis-inducing conditions (e.g., GPX4 inhibition by RSL3 or ML210), the accumulation of lipid hydroperoxides (PLOOH) leads to membrane rupture. **b**, Immunoblot (IB) analysis of HMOX1, and vinculin in A375 sgNT and HMOX1^KO^ cells. **c**, Dose-dependent cytotoxicity in A375 sgNT and HMOX1^KO^ cells treated with increased concentration of hemin (0 – 128 μM) for 72 h. **d**, Dose-dependent cytotoxicity in A375 sgNT and HMOX1^KO^ cells pre-treated with hemin for 24 h (10 μM) followed by ML210 treatment for 72 h. **e**, Immunoblot (IB) analysis of HMOX1 and vinculin in A375 HMOX1^KO^ cells reconstituted with either an empty vector control or a vector encoding HMOX1. **f**, Dose-dependent cytotoxicity of 10 μM hemin (24 h) and ML210 (72 h) treatment in A375 sgNT and A375 HMOX1^KO^ cells reconstituted with either an empty vector control or a vector encoding HMOX1. **g**, Crystal violet staining of A375 HMOX1^KO^ cells reconstituted with either an empty vector control or a vector encoding HMOX1 after 10 μM hemin and 0.1 μM ML210 co-treatment. **h**, Dose-dependent cytotoxicity in A375 HMOX1^KO^ cells reconstituted with either an empty vector control or a vector encoding HMOX1, after hemin pre-treatment (10 μM) followed by ML210, Erastin or H2O2 (72 h). Cell viability was assessed using Alamar Blue. Data represent mean ± SD of triplicates from one representative experiment out of three performed.

To confirm that the protective effect was dependent on HMOX1 deficiency, we tested whether the HMOX1 knockout in A375 HMOX1^KO^ cells could be complemented by transducing the cells with lentiviral vectors carrying either wild-type HMOX1 or an empty vector cassette (control) (Fig. 1e). To validate these cells, we treated them with an increasing concentration of hemin. As expected, reintroduction of HMOX1 rendered the A375 HMOX1^KO^ cells (HMOX1-reconstituted cells) less sensitive to high doses of hemin (Extended Data 1e). These cells also regained sensitivity to co-treatment with 10 μM hemin and ML210, as assessed by Alamar Blue and crystal violet cell viability staining. In contrast, cells transduced with an empty lentiviral vector became more resistant to ferroptosis after 10 μM hemin and ML210 co-treatment (Fig. 1f,g). Notably, hemin protected HMOX1^KO^ cells transduced with an empty lentiviral vector against GPX4i, but it did not confer protection against Erastin, hydrogen peroxide, or other cytotoxic agents tested with various mechanisms of action (Fig. 1h, Extended Data 1f).

### 2. Non-toxic hemin supplementation protects HMOX1/GPX4 double-knockout cells from ferroptosis

Since accumulation of lipid peroxides is a hallmark of ferroptosis, we monitored lipid peroxidation using the BODIPY-C11 probe. The BODIPY-C11 probe is a fluorescent dye that shifts from red to green emission upon oxidation, allowing it to serve as a reporter of lipid peroxidation in membranes. We observed that HMOX1 deficiency attenuated lipid peroxidation triggered by combined hemin and ML210 treatment (Extended Data 2a). In addition, because lipid metabolism is a key feature in ferroptosis, we explored the lipidomic profile of cells treated with hemin. This analysis showed no significant difference in the levels of polyunsaturated species across various phospholipid classes (Extended Data 2b-e).

Because both pharmacological inhibition (ML210) and genetic knockout of GPX4 trigger ferroptosis, we tested whether hemin could protect cells in each context, thereby excluding off-target effects or residual GPX4 activity. To do this, we knocked out GPX4 in A375 wild-type cells, generating GPX4^KO^, and in A375 HMOX1^KO^ cells, generating GPX4^KO^/HMOX1^KO^ (Fig. 2a). We found that disrupting HMOX1 in GPX4^KO^ cells increased cell viability by approximately 70%, an effect likely mediated by increased heme availability (Fig. 2b). This finding was consistent with the protection of GPX4^KO^/HMOX1^KO^ hemin-treated cells observed by microscopy (Extended Data 2f). To confirm that the protective effect required HMOX1 absence, we reconstituted HMOX1 expression in GPX4^KO^/HMOX1^KO^ cells using a doxycycline-inducible vector, which restored HMOX1 after doxycycline treatment and abolished hemin-mediated protection (Fig. 2c,d).

**Fig. 2.**
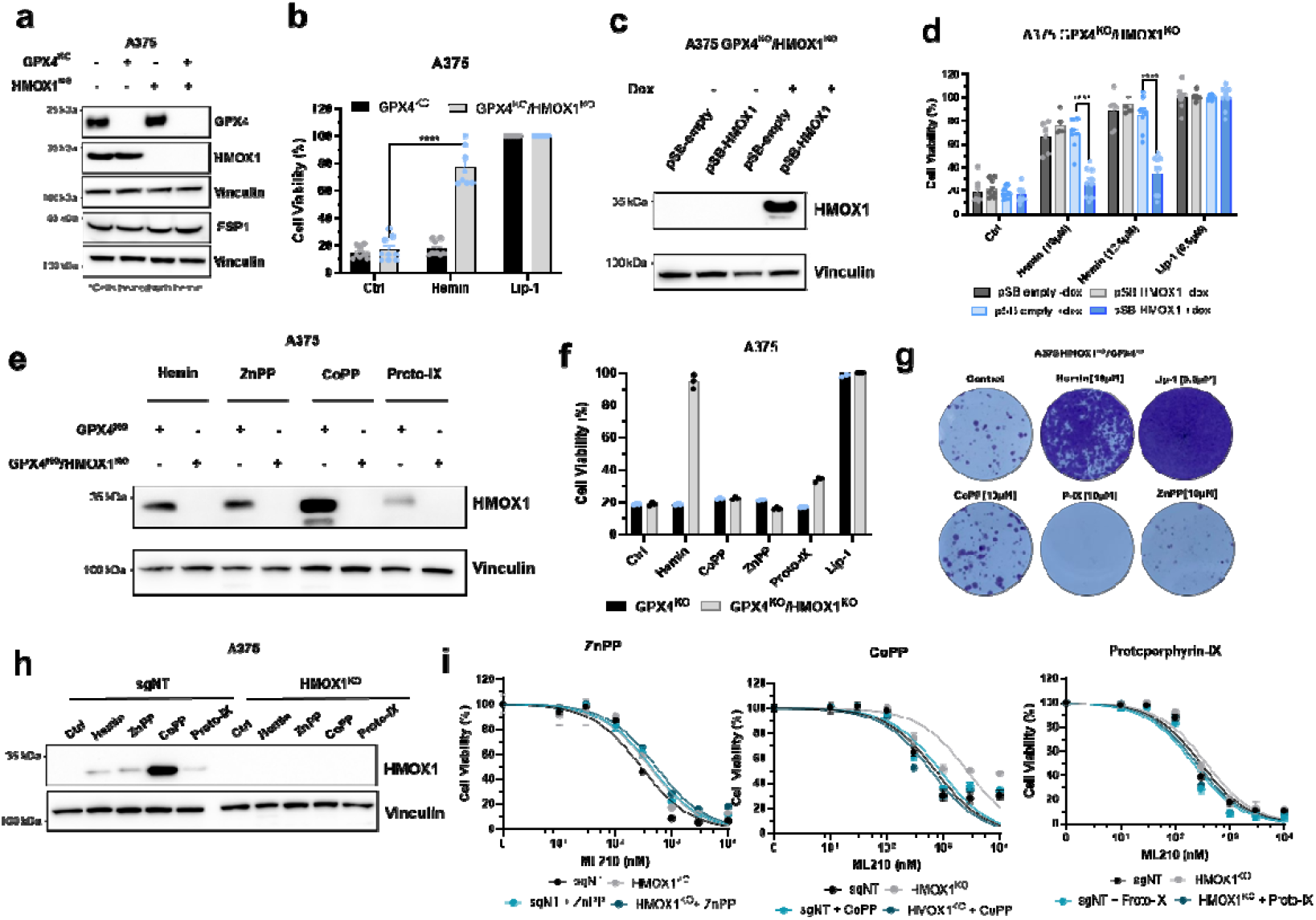
Non-toxic hemin supplementation protects HMOX1/GPX4 double-knockout cells from ferroptosis. **a**, Immunoblot (IB) analysis of GPX4, HMOX1, FSP1 and vinculin in A375 GPX4^KO^ and GPX4^KO^/HMOX1^KO^ cells. **b**, Cell viability of A375 GPX4^KO^ and GPX4^KO^/HMOX1^KO^ after treatment with hemin (10 μM, 72 h) or lip-1 (500 nM, 72h). Cell viability was assessed using Alamar Blue. Data represent mean ± SD of triplicates from three independent experiments. *P□<□0.05, * *P□<□0.01, * * *P□<□0.001, * * * *P□<□0.0001. Each dot represents a replicate. **c**, Immunoblot (IB) analysis of GPX4, HMOX1 and vinculin in A375 GPX4^KO^/HMOX1^KO^ cells reconstituted with either the pSB empty vector or the pSB vector encoding HMOX1. **d**, Cell viability of A375 GPX4^KO^/HMOX1^KO^ cells reconstituted with either the pSB empty vector or the pSB vector encoding HMOX1 after treatment with hemin (10 μM, 72 h) or lip-1 (500 nM, 72 h). Cell viability was assessed using Alamar Blue. Data represent mean ± SD of triplicates from three independent experiments. *PL<□0.05, * *P□<□0.01, * * *P□<□0.001, * * * *P□<□0.0001. Each dot represents a replicate. **e**, Immunoblot (IB) analysis of HMOX1 in A375 GPX4^KO^ and GPX4^KO^/HMOX1^KO^ cells treated with hemin, ZnPP, CoPP, or Protoporphyrin IX (10 μM, 24 h). **f**, Cell viability of A375 GPX4^KO^ and GPX4^KO^/HMOX1^KO^ cells after treatment with hemin, ZnPP, CoPP, or Protoporphyrin IX (10 μM, 24 h) or lip-1 (500 nM, 72h). Cell viability was assessed using Alamar Blue. Data represent mean ± SD of triplicates from three independent experiments. *P□<□0.05, * *P□<□0.01, * * *P□<□0.001, * * * *P□<□0.0001. Each dot represents a replicate. **g**, Crystal violet staining of A375 GPX4^KO^ and GPX4^KO^/HMOX1^KO^ cells after treatment with hemin, ZnPP, CoPP, Protoporphyrin IX (10 μM, 24 h) or lip-1 (500 nM, 72h). **h**, Immunoblot (IB) analysis of HMOX1 and vinculin expression in A375 sgNT and HMOX1^KO^ cells. **i**, Dose-dependent cytotoxicity of 10 μM ZnPP, CoPP, or Protoporphyrin IX in A375 sgNT and HMOX1^KO^ cells pre-treated with the analogs at 24 h followed by ML210 treatment at 72 h. Cell viability was measured using Alamar Blue. Data represent mean ± SD of triplicates from one representative experiment out of two.

To determine whether this effect was specifically triggered by hemin (protoporphyrin ring + iron) rather than the protoporphyrin ring alone, we treated HMOX1^KO^ and GPX4^KO^/HMOX1^KO^ cells with protoporphyrin analogs ZnPP, CoPP, and Proto-IX. All analogs induced HMOX1 expression, but did not affect cell viability after pharmacological inhibition of GPX4 with ML210 (GPX4i) or in GPX4^KO^ cells (Fig. 2e-i). Collectively, these findings demonstrate that hemin exerts a context-dependent and paradoxical function in ferroptosis: in the presence of HMOX1, it enhances GPX4i-or GPX4^KO^-induced cell death, whereas in its absence, it unexpectedly protects cells by increasing their resistance to ferroptosis.

### 3. Catalytic inactivation of HMOX1, not nuclear localization, underlies heme-mediated ferroptosis resistance

Previous studies have shown that HMOX1 is anchored to the endoplasmic reticulum (ER), where it catalyzes heme degradation ^20^. However, under stress, HMOX1 can be cleaved from the ER and translocated to the nucleus to act in an enzymatic-independent manner ^21^. Given the dual and context-dependent role of heme in ferroptosis, acting to either increase or decrease cellular sensitivity, we investigated whether this role was dependent on the catalytic activity or cellular localization of HMOX1. To do this, we used a truncated form of the protein shown to localize to the nucleus. Specifically, we reconstituted HMOX1^KO^ cells with HMOX1 variant truncated at serine 275, referred to as HMOX1^trunc 21,22^. It localized to the nucleus, as confirmed by immunofluorescence (Fig. 3a). These cells exhibited increased resistance to hemin toxicity, confirming successful reintroduction of the protein (Fig. 3b).

**Fig. 3.**
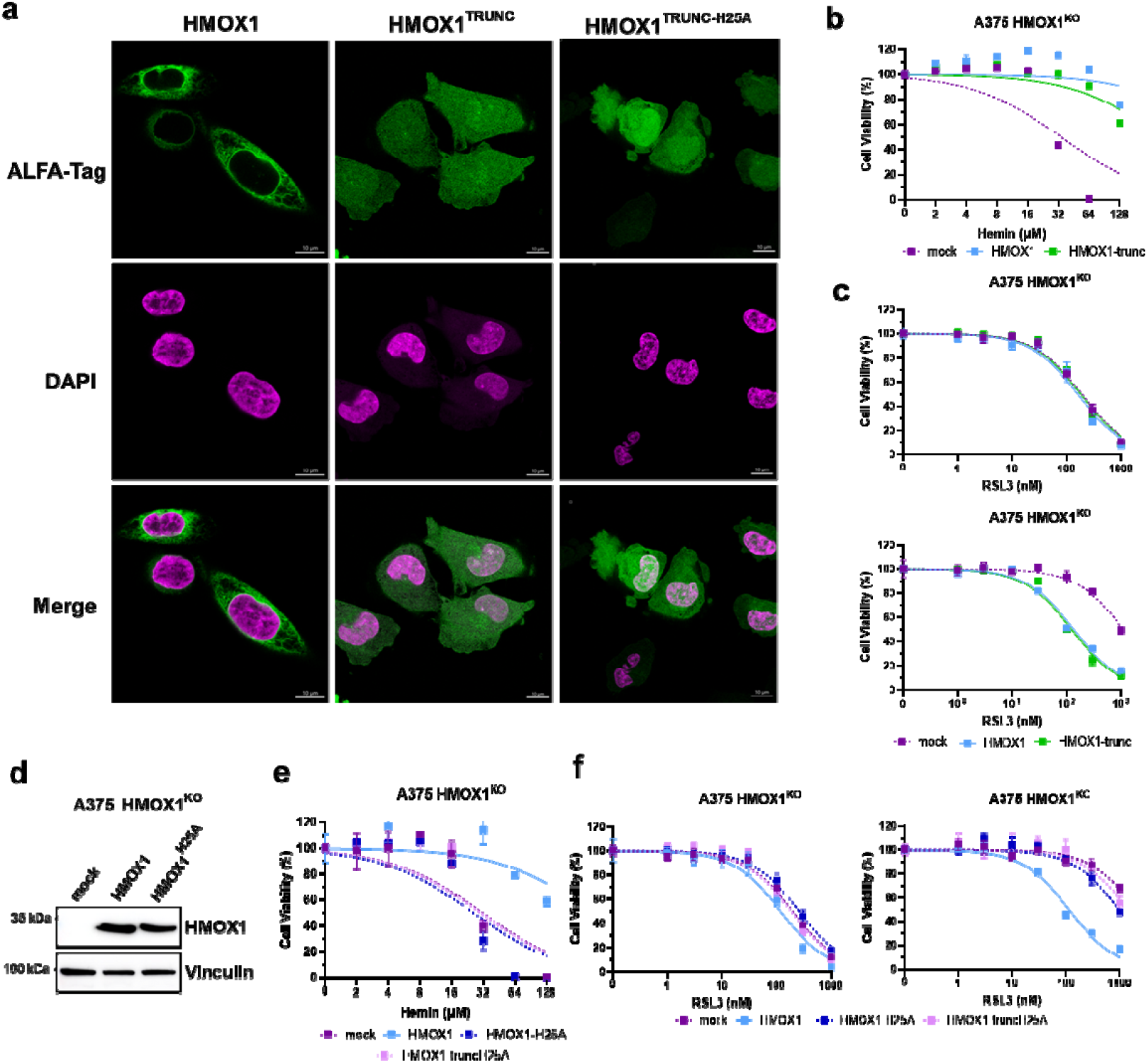
Catalytic inactivation of HMOX1, not nuclear localization, underlies heme-mediated ferroptosis resistance. **a**, Immunofluorescence analysis of A375 HMOX1^KO^ cells reconstituted with either an empty vector control or a vector encoding either HMOX1, HMOX1-trunc, or HMOX1-truncH25A tagged with an alfa-tag. Scale bar, 10 μm. **b**, Dose-dependent cytotoxicity of increased concentration of hemin (0 – 128 μM, 72 h) in A375 HMOX1^KO^ cells reconstituted with either an empty vector control or a vector encoding either HMOX1, or HMOX1-trunc. **c**, Dose-dependent cytotoxicity of 10 μM hemin (24 h) and RSL3 (72 h) in A375 HMOX1^KO^ cells reconstituted with either an empty vector control or a vector encoding either HMOX1, or HMOX1-trunc. **d**, Immunoblot (IB) analysis of HMOX1, and vinculin in A375 HMOX1^KO^ cells reconstituted with either an empty vector control or a vector encoding HMOX1, or HMOX1-H25A. **e**, Dose-dependent cytotoxicity of increased concentration of hemin (0 – 128 μM, 72 h) in A375 HMOX1KO cells reconstituted with either an empty vector control or a vector encoding either HMOX1, HMOX1-H25A, or HMOX1-truncH25A. **f**, Dose-dependent cytotoxicity of 10 μM hemin (24 h) and RSL3 (72 h) in A375 HMOX1^KO^ cells reconstituted with either an empty vector control or a vector encoding HMOX1, HMOX1-H25A, or HMOX1-truncH25A. Cell viability was assessed using Alamar Blue. Data represent mean ± SD of triplicates from a representative experiment, repeated three times independently.

Interestingly, ferroptosis-resistant HMOX1^KO^ cells reconstituted with HMOX1^trunc^ were re-sensitized to GPX4 inhibition (Fig. 3c). This indicated that although HMOX1 can translocate to the nucleus, its concurrent cytosolic presence ensures that nuclear localization does not contribute to the heme-mediated ferroptosis resistance. We therefore tested whether loss of HMOX1 catalytic activity, independent of nuclear localization, was required for resistance. To do this, we reconstituted HMOX1^KO^ cells with a catalytically inactive mutant, HMOX1^H25A 22^, either wild-type or truncated (HMOX1^trunc-H25A^) (Fig. 3d). These cells were more sensitive to toxic doses of hemin, confirming loss of enzymatic activity (Fig. 3e). In addition, both HMOX1^H25A^-and HMOX1^trunc-H25A^-reconstituted cells retained their resistance to non-toxic hemin treatment (Fig. 3f). These findings suggest that the absence of HMOX1 catalytic activity likely leads to heme accumulation, which is required to protect cells from ferroptosis.

### 4. Heme stabilization drives ferroptosis resistance independently of the BACH1– NRF2 pathway

Heme promotes proteasomal degradation of the transcriptional repressor BACH1, thereby activating the transcriptional factor NRF2 and inducing a network of antioxidant genes, including HMOX1, that can protect cells from ferroptosis ^23^. Thus, we hypothesized that in the absence of HMOX1, intracellular heme would be expected to accumulate, thereby promoting more efficient BACH1 degradation and NRF2 activation and leading to ferroptosis resistance ^24^. As expected, hemin supplementation reduced BACH1 protein levels (Extended Data 3a). To directly assess the role of BACH1 in this context, we generated BACH1-deficient cell lines (Extended Data 3b). As shown in Figure 4a, BACH1 knockout alone was not sufficient to protect cells from ferroptosis. Increased resistance to GPX4i was observed only in HMOX1^KO^ cells treated with hemin (Fig. 4a). Next, to test whether NRF2 contributes to ferroptosis protection, we disrupted HMOX1 in NRF2-deficient cells (NRF2^KO^) and re-expressed wild-type NRF2 across all cell lines. We validated successful generation of these cell lines by elevated expression of NRF2 target genes, such as xCT, FSP1, and NQO1 (Extended Data 4c). As shown in Figure 4b, disruption of HMOX1 in NRF2^KO^ protected these cells from ferroptosis, likely through the accumulation of heme. Altogether, these data indicate that heme-mediated ferroptosis resistance in HMOX1^KO^ cells is BACH1 and NRF2-independent.

**Fig. 4.**
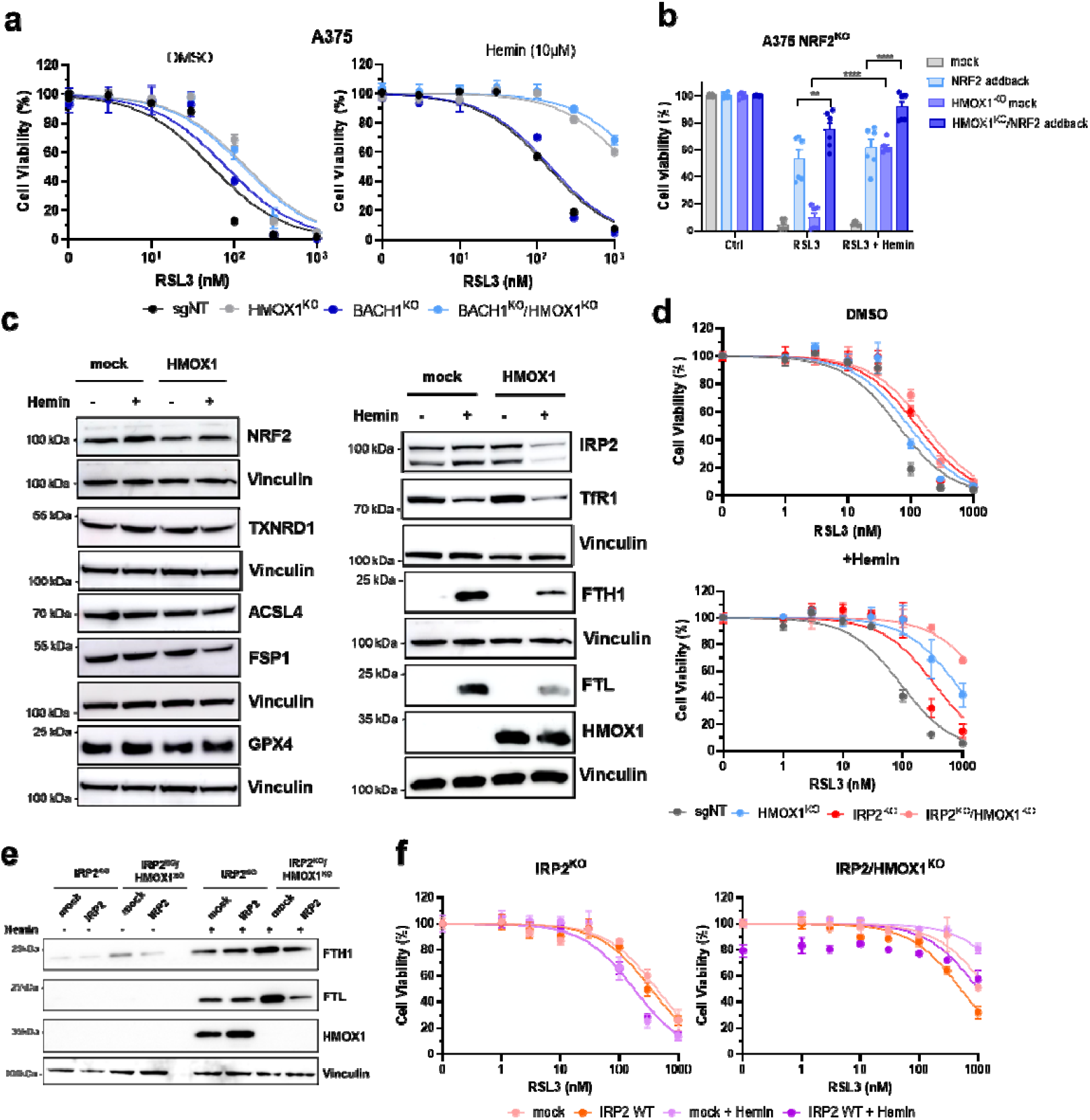
Heme stabilization drives resistance independently of the BACH1–NRF2 pathway: **a**, Dose-dependent cytotoxicity of 10 μM hemin pre-treatment (24 h) and ML210 treatment (72 h) in A375 sgNT, HMOX1^KO^, BACH1^KO^ and HMOX1^KO^/BACH1^KO^ cells. **b**, Cell viability of A375 NRF2^KO^ and NRF2^KO^/HMOX1^KO^ cells reconstituted with NRF2 after hemin pre-treatment (10 μM, 24 h), followed by RSL3 (72 h). Cell viability was assessed using Alamar Blue. Data represent mean ± SD of triplicates from three independent experiments. *P□<□0.05, * *P□<□0.01, * * *P□<□0.001, * * * *P□<□0.0001. Each dot represents a replicate. **c**, Immunoblot (IB) analysis of NRF2, TXNRD1, ACSL4, FSP1, GPX4, IRP2, TfR1, FTH1, FTL, HMOX1, and vinculin in A375 HMOX1^KO^ cells reconstituted with an empty vector control or a vector encoding HMOX1. **d**, Dose-dependent cytotoxicity of 10 μM hemin pre-treatment (24 h) and ML210 treatment (72 h) in A375 sgNT, HMOX1^KO^, IRP2^KO^ and HMOX1^KO^/IRP2^KO^ cells. **e**, Immunoblot (IB) analysis of FTH1, FTL, HMOX1 and vinculin in A375 HMOX1^KO^ and HMOX1^KO^/IRP2^KO^ reconstituted with an empty vector control or a vector encoding IRP2. **f**, Dose-dependent cytotoxicity of 10 μM hemin pre-treatment (24 h) and RSL3 treatment (72 h) in A375 HMOX1^KO^ and HMOX1^KO^/IRP2^KO^ reconstituted with an empty vector control or a vector encoding IRP2. Cell viability was measured using Alamar Blue. Data represent mean ± SD of triplicates from one representative of three independent experiments.

The transcription factor NRF2 induces the expression of ferritin, the major intracellular iron storage complex ^25^. Ferritin is composed of heavy (ferritin heavy chain 1, FTH1) and light (ferritin light chain, FTL) chain subunits. By sequestering labile iron, ferritin prevents iron-dependent lipid peroxidation and thereby protects cells from ferroptosis ^26^. Surprisingly, we observed that FTH1 was strongly induced by hemin even in the absence of NRF2 (Extended Data 4d,e). We further examined the expression of a broad range of ferroptosis-related genes after hemin treatment and found that only FTH1 and FTL were differentially induced (Fig. 4c). In contrast to NRF2, which promotes ferritin expression, the iron regulatory protein IRP2 post-transcriptionally represses FTH1 and FTL, thereby limiting ferritin translation and increasing ferroptosis sensitivity ^27,28^. Interestingly, although IRP2 levels were not reduced in HMOX1^KO^ cells after hemin treatment, its activity appeared diminished, as its targets FTH1 and FTL were induced while TfR1 was suppressed (Fig. 4c). To investigate whether IRP2 contributes to ferritin regulation in the absence of HMOX1, we generated IRP2 knockout and IRP2/HMOX1 double knockout cells. We observed that loss of IRP2 increased expression of FTHL and FTL, and this effect was enhanced by HMOX1 absence (Extended Data 3f). Consistently, these cells were more resistant to ferroptosis (Fig. 4d). Notably, hemin treatment still induced ferritin expression and conferred ferroptosis resistance even in IRP2^KO^ and IRP2^KO^/HMOX1^KO^ cells (Fig. 4d, Extended Data 3g). Using IRP2-reconstituted cells (IRP2 add-back), we confirmed that lack of IRP2 contributes to ferritin upregulation in the absence of HMOX1, as IRP2-reconstituted cells had lower ferritin levels (Fig. 4e). Furthermore, in IRP2-reconstituted cells, hemin treatment provided reduced protection (Fig 4f), indicating that decreased ferritin levels may attenuate heme-mediated resistance. Collectively, these findings suggest that heme, likely stabilized in the absence of HMOX1, suppresses ferroptosis independently of BACH1 and NRF2, while IRP2 inactivation partially contributes to ferritin upregulation and resistance to ferroptosis.

### 5. Heme stabilization prevents ferroptosis by regulating ferritin levels

Building on our previous observations, we hypothesized that ferritin may contribute to hemin-mediated ferroptosis resistance. To investigate this, we examined the levels of FTH1 and FTL in HMOX1^KO^ cells transduced with a vector encoding either a control cassette, wild-type HMOX1, or a catalytically inactive mutant, HMOX1^H25A^. As depicted in Figure 5a, supplementation with hemin increased FTH1 and FTL levels in both HMOX1^KO^ and HMOX1^H25A^-reconstituted HMOX1^KO^ cells, but not in HMOX1-reconstituted HMOX1^KO^ cells. Notably, hemin was the only protoporphyrin able to induce FTH1 and FTL in HMOX1^KO^ cells (Fig. 5b). Because FTH1 contains the ferroxidase site required for iron oxidation, which limits the labile iron pool that drives lipid peroxidation during ferroptosis, we generated FTH1 knockout (FTH1^KO^) and FTH1/HMOX1 double-knockout (FTH1^KO^/HMOX1^KO^) cells (Fig. 5c). Notably, hemin failed to confer ferroptosis protection in the FTH1^KO^/HMOX1^KO^ cells (Fig. 5d). In addition, FTH1^KO^ cells were sensitive to hemin, with the effect most pronounced in double knockout cells, indicating a strong dependence on ferritin under these conditions (Extended Data 4a-c). Next, we reintroduced FTH1 in FTH1^KO^ and FTH1^KO^/HMOX1^KO^ cells (Fig. 5e), which rescued heme-mediated ferroptosis protection (Fig. 5f). These findings indicate that heme, likely stabilized and accumulated in HMOX1-deficient cells, mediates ferroptosis suppression partly by upregulating ferritin.

**Fig. 5.**
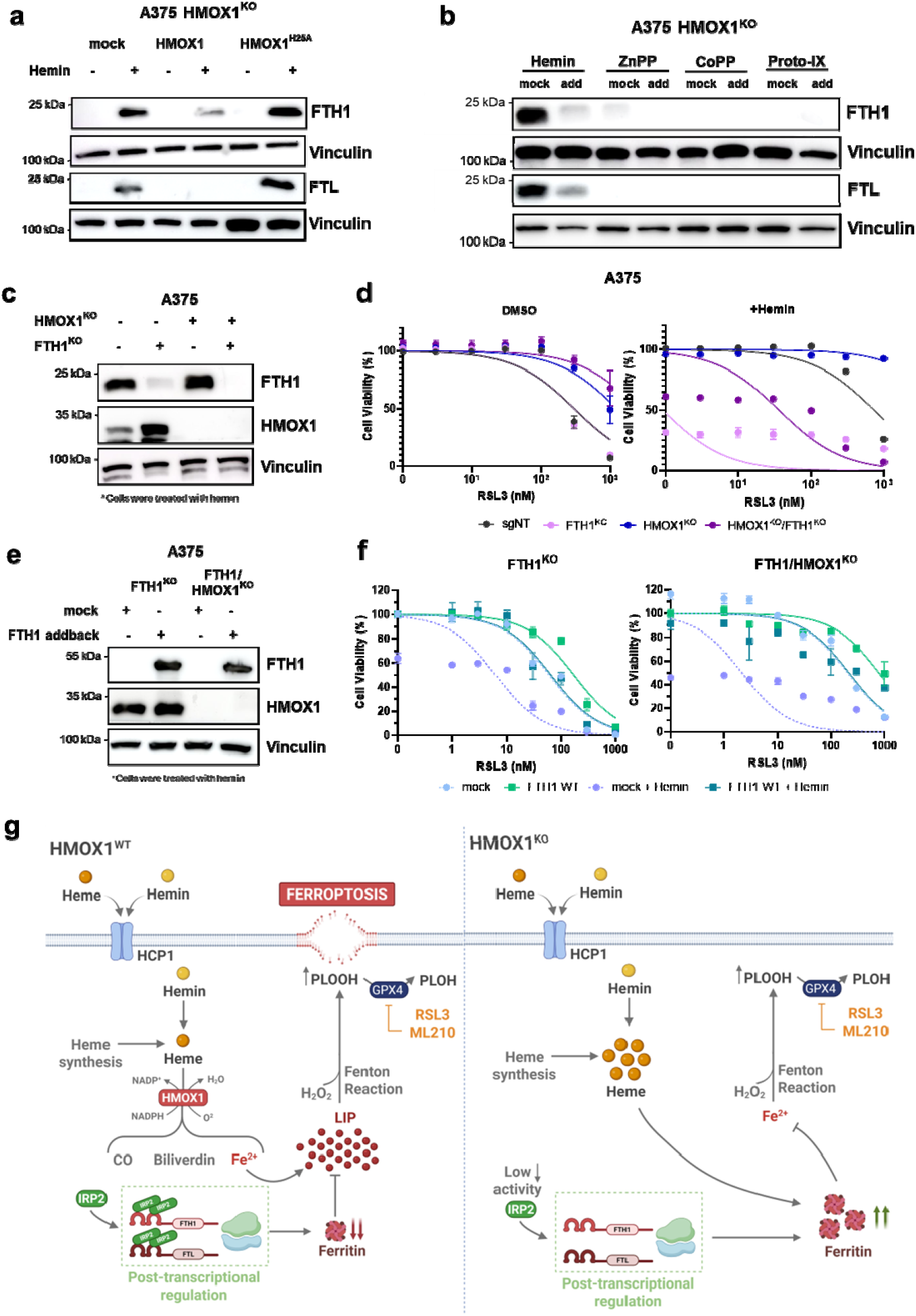
Heme stabilization prevents ferroptosis by regulating ferritin levels. **a**, Immunoblot (IB) analysis of FTH1, FTL, and vinculin in A375 HMOX1^KO^ cells reconstituted with an empty vector control or a vector encoding HMOX1 or HMOX1^H25A^. **b**, Immunoblot (IB) analysis of FTH1, FTL, and vinculin in A375 HMOX1^KO^ cells reconstituted with an empty vector control or a vector encoding HMOX1 after treatment with hemin, ZnPP, CoPP or Protoporphyrin-IX (10 μM, 24 h). **c**, Immunoblot (IB) analysis of FTH1, HMOX1, and vinculin in A375 sgNT, HMOX1^KO^, FTH1^KO^ and HMOX1^KO^/FTH1^KO^ cells. **d**, Dose-dependent cytotoxicity of 10 μM hemin pre-treatment (24 h) and RSL3 treatment (72 h) in A375 sgNT, HMOX1^KO^, FTH1^KO^ and HMOX1^KO^/FTH1^KO^ cells. Cell viability was measured using Alamar Blue. Data represent mean ± SD of triplicates from one representative of three independent experiments. **e**, Immunoblot (IB) analysis of FTH1, HMOX1, and vinculin in A375 FTH1^KO^ and HMOX1^KO^/FTH1^KO^ cells reconstituted with either an empty vector control or a vector encoding FTH1. **f**, Dose-dependent cytotoxicity of 10 μM hemin pre-treatment (12 h) and RSL3 treatment (72 h) in A375 FTH1^KO^ and HMOX1^KO^/FTH1^KO^ cells reconstituted with either an empty vector control or a vector encoding FTH1. **g**, Schematic view of the uncovered role of heme in ferroptosis. Once in the cytosol, heme can be catabolized by heme oxygenase-1 (HMOX1), generating biliverdin, carbon monoxide (CO), and ferrous iron (Fe2□). The liberated iron feeds the labile iron pool (LIP), which drives lipid peroxidation and triggers ferroptotic cell death. Under ferroptosis-inducing conditions, such as GPX4 inhibition by RSL3 or ML210, lipid hydroperoxides (PLOOH) accumulate, ultimately causing plasma membrane disruption. When HMOX1 activity is absent, heme accumulates and induces ferritin expression, which sequesters free iron, thereby limiting LIP expansion and suppressing ferroptosis. Cell viability was measured using Alamar Blue. Data represent mean ± SD of triplicates from one representative of four independent experiments.

## Discussion

Beyond its role as an iron donor, the intact heme molecule exerts regulatory effects on signaling and drug metabolism, but these functions in cancer biology and ferroptosis remain unclear ^29^. Intracellular heme accumulation activates HMOX1, generating metabolites with antioxidant and anti-inflammatory properties. At the same time, excess heme expands the pool of chelatable, redox-active iron, thereby promoting ferroptosis ^30^. Despite these observations, how heme signaling intersects with ferroptosis, particularly in cancer, remains poorly understood. Here, we uncover a previously unknown role of heme in ferroptosis, showing that it functions as an HMOX1-dependent switch that determines whether cells undergo or resist ferroptosis. This dual effect depends on the catalytic activity of HMOX1: in its absence, heme stabilizes and elevates ferritin levels, conferring marked protection against GPX4 inhibition (Fig. 5g).

Although heme can be obtained from dietary sources, it is primarily synthesized endogenously through a multi-step process involving both mitochondrial and cytosolic compartments ^30^. Emerging evidence shows that cancer cells have increased levels of heme and its biosynthetic precursors, higher activity of heme-containing enzymes, and greater regulation of heme import and export channels ^31,32^. These observations suggest that tumors actively reprogram heme metabolism, beyond iron levels, to support malignant proliferation and survival. While increased heme synthesis and import contribute to its intracellular accumulation, its steady-state concentration is primarily regulated by HMOX1-mediated degradation ^33,34^. Previous studies have shown that increased heme degradation via HMOX1 can suppress tumor growth through ferroptosis ^12,35-37^. In this study, we confirmed this observation by showing that increased HMOX1 levels induced by hemin promote ferroptosis sensitivity in various cancer cell types. However, we also found that in the absence of HMOX1, heme exerts a strong antiferroptotic activity. Specifically, loss of HMOX1 enzymatic activity leads to intracellular heme accumulation, which inhibits ferroptotic cell death.

Previous studies have suggested that HMOX1 deficiency leads to cellular heme toxicity, whereas inactive HMOX1 mutants can paradoxically confer protection against oxidative stress through mechanisms that remain only partially understood ^22,38-41^. To address this gap, we identified ferritin as a central mediator of heme-mediated ferroptosis resistance in HMOX1-deficient cells. FTH1 knockout cells were hypersensitive to ferroptosis upon hemin treatment, whereas reintroduction of FTH1 fully restored resistance. These findings are consistent with a role for ferritin in buffering intracellular iron and protecting against ferroptotic cell death ^7,42-44^.

The role of heme in cancer has traditionally been examined in the context of dietary exposure, particularly from red and processed meats, which contain heme as hemoglobin and myoglobin ^30^. Studies have consistently linked high consumption of these foods to increased incidence of several carcinomas, for example, colorectal cancer ^45^. Accordingly, the International Agency for Research on Cancer (IARC) has classified processed meat as carcinogenic to humans (Group 1) and red meat as probably carcinogenic (Group 2A), implicating dietary heme as a potential key contributor to tumorigenesis ^46^. In addition, hemin is therapeutically administered for acute porphyrias, highlighting the biological significance of exogenous heme in disease ^47^. Our study expands this perspective by showing that exogenously acquired heme, when stabilized by loss of HMOX1 activity, promotes ferroptosis resistance across multiple tumor types, including melanoma, lung carcinoma, and fibrosarcoma. These findings suggest that heme may support tumor survival by protecting cells from regulated cell death, revealing a previously unrecognized facet of heme’s involvement in cancer biology.

## Materials and Methods

### Chemicals and reagents

Hemin (cat. no. H9039), Liproxstatin-1 (SML1414), ML210 (SML0521), RSL3 (SML2234), ZnPP (691550-M), CoPP (C1900), and Proto-IX (P8293) were obtained from Sigma-Aldrich. BODIPY-C11 (cat. no. D3861) was purchased from Thermo Fisher Scientific. KI969 was purchased from MedChem Express (cat. no. HY-101140). All drugs were prepared as stock solutions in DMSO under sterile conditions. Stock solutions were stored at -20 °C.

### Cell culture conditions

The cell lines A375 (human melanoma), A549 (human lung carcinoma), and HT1080 (human fibrosarcoma) were purchased from ATCC. All cell lines were authenticated by STR profiling and verified to be negative for mycoplasma contamination. They were cultured in Dulbecco’s modified Eagle’s medium (DMEM; Gibco), supplemented with 10% fetal bovine serum (FBS; Gibco) and 100 U/mL penicillin/streptomycin (P/S; Gibco), at 37□°C in a humidified 5% CO_2_ atmosphere.

### Cell viability assays

For the Alamar Blue assay, 3,000 cells were seeded on 96-well plates and pre-treated with hemin, ZnPP, CoPP, or Proto-IX for 24 h (unless stated otherwise). Next, RSL3 or ML210 were added and incubated with the cells for 72 h. Cell viability was assessed using Alamar Blue. Alamar Blue solution was prepared by dissolving 1 g of resazurin sodium salt in 200 mL of sterile PBS and filtered through a 0.22 µm filter. Stock solutions were stored at 4°C. The working solution was made freshly by adding 100 µL of the stock solution to 10mL of growth media. After 1-2h incubation time, viability was estimated by measuring the fluorescence using a 540/35 excitation filter and a 590/20 emission on a Spark® microplate reader (Tecan, Zürich, Switzerland).

For the clonogenic assay, 5,000 cells were seeded per well in a 12-well plate. Hemin, ZnPP, CoPP, or Proto-IX were added 6 h and RSL3 or ML210 24 h after seeding. The plates were cultured for 10 days under respective conditions. The colonies were stained with 0.01% (w/v) crystal violet-PBS solution.

### Generation of knockout cell lines

Single sgRNA guides were chosen using the VBC score (https://www.vbc-score.org/)53. Guide sequences were cloned into the pLenti-CRISPR-V2 vector by annealing oligonucleotides (Eurofins genomics) with specific overhangs complementary to BsmBI-digested sites (Addgene #98293 and #98290). Plasmid DNA was confirmed by Sanger sequencing. Cells were transduced with lentiviruses expressing the indicated constructs and subjected to selection for 7 days. Cells were used as pools, or single clones were generated when necessary. Loss of protein expression in cells was validated by Western blot analysis. The following single-guide RNA (sgRNA) primer sets were used:

hHMOX1_sgRNA3 – TACCTGGGTGACCTGTCTGG

hGPX4_sgRNA1 – GAAGCGCTACGGACCCATGG

hBACH1_sgRNA1 – GTACCTGCGGAGTTTACACTG

hNRF2_sgRNA1 – GGATCTGCCAACTACTCCC

hIRP2_sgRNA2 – AAAATCTTGAAGAAGAACAC

hFTH1_sgRNA1 – GCACCATGGACAGGTAAACGT

Lentiviral particles were produced with a third-generation packaging system and X-tremeGENE HP DNA Transfection Reagent (Sigma-Aldrich, 6366546001). HEK293T cells were transfected with the lentiviral transfer plasmids, two packaging plasmids (pMDLg/ pRRE, Addgene #12251 and pRSV-Rev, Addgene #12253), and the envelope plasmid (pCMV-VSV-G, Addgene#8454). Viral particles present in the cell culture supernatant were harvested 48 and 72 h after transfection and used to transduce the target cell line by directly incubating cells with HEK293T supernatants filtered through a 0.44µM membrane. All experimental procedures for lentivirus production and transduction were performed in a biosafety level 2 laboratory.

### Lentivirus production for gene overexpression

Codon-optimized HMOX1 and IRP2 were synthesized by IDT as gBlocksTM and subcloned into the p442-IRES vector. The following constructs were generated: p442-IRES-Mock, p442-IRES-HMOX1-Alfatag, p442-IRES-HMOX1(trunc)-Alfatag, p442-IRES-HMOX1(trunc-H25A)-Alfatag, p442-IRES-HMOX1(H25A)-Alfatag, and p442-IRES-IRP2-Flag. HMOX1 overexpression constructs were additionally generated using the transposase vector pSB-ET-iE, as previously described (doi: 10.1038/s41388-020-01477-8). pSB-based expression vectors were co-transfected with the Sleeping Beauty transposase vector pCMV-(CAT)T7-SB100, generating pSB-HMOX1. The cDNA of NFE2L2 was subcloned from pSB-NFE2L2 into p442-IRES, generating the p442-IRES-NFE2L2 plasmid. Similarly, the cDNA of FTH1 was subcloned into p442-IRES and fused to FKBP12^F36V^. All plasmids were verified by Sanger sequencing and subsequently used for lentiviral transduction.

### GPX4^KO^ cell experiments

A375 cells deficient in GPX4 or harboring double knockouts were cultured in medium supplemented with 0.5 μM Lip-1 to inhibit ferroptosis. One day after plating, the medium was removed and cells were washed once with 200 μL of 1× PBS. Following the wash, cells were incubated with medium containing the indicated treatments of interest with or without Lip-1 supplementation. Cell death was determined 72 h later.

### Western blot

Cell lysates were prepared using Radioimmunoprecipitation assay (RIPA) buffer supplemented with a protease inhibitor cocktail (Roche). Proteins were separated by SDS– polyacrylamide gel electrophoresis and electrotransferred onto polyvinylidene difluoride membranes. Membranes were subsequently blocked with 5% non-fat milk in Tris-buffered saline containing 0.1 % Tween 20 (TBS-T) for 1 h at room temperature (RT). Primary antibodies were diluted in 5% bovine serum albumin (BSA) in TBS-T solution, and membranes were incubated with the primary antibody solution at 4°C overnight. After washing with TBS-T, membranes were incubated at RT for 2 h with horseradish peroxidase-conjugated secondary antibody (1:3,000, anti-rabbit #7074, anti-mouse #7076, or anti-rat) diluted in 5% BSA in TBS-T. Protein bands were visualized using the Immobilon Western Chemiluminescent HRP Substrate (Sigma-Aldrich) and detected by enhanced chemiluminescence. Images were acquired using the Amersham ImageQuant 800 detection system (Cytiva). The following antibodies were used against Vinculin: 1:1000, sc-73614, Santa Cruz Biotechnology; HMOX1: 1:1000, D60G11, Cell Signaling; GPX4: 1:1,000, ab125066, Abcam; FSP1: 1:10, antibody raised against recombinant human FSP1 protein, clone 6D8-11, developed in M. Conrad (Munich, Germany); xCT: 1:10, antibody raised against recombinant human xCT protein, developed in M. Conrad (Munich, Germany); IRP2: 1:1000, D6E6W, Cell Signaling; FTH1: 1:1,000, ab65080, Abcam; FTL: 1:1,000, ab69090, Abcam; TfR1: 1:1,000, ab84036, Abcam; NRF2: 1:1,000, ab62352, Abcam; TXNRD1: 1:1000, D1T3D, Cell Signaling; β-actin: 1:5,000, A5441, Sigma-Aldrich; BACH1: 1:1000, sc-271211, Santa Cruz; ACSL4: 1:1000, ab205197, Abcam; FluoTag®-X2 anti-ALFA: 1:500 for IF, 1:1000 for WB, N1502-At488-L, NanoTag Biotechnologies.

### Assessment of lipid peroxidation using C11-BODIPY (581/591)

150,000 cells per well were seeded in 6-well plates one day before the experiment. The next day, cells were treated for 3 h and then incubated with C11-BODIPY (581/591) (1□μM) for 30□min at 37□°C. Subsequently, cells were trypsinized, resuspended in 500□μL of fresh PBS, and analyzed by flow cytometry using a 488-nm excitation laser to record fluorescence (FACS Canto II, BD Biosciences) in PE and FITC channels. The samples were analyzed using FlowJo software (BD Biosciences). Cells were first gated based on forward and side scatter (FSC-A vs SSC-A) to exclude debris. Viable cells were selected based on scatter properties. The C11-BODIPY fluorescence intensity (FITC channel, 488 nm excitation/510–550 nm emission) was quantified within the viable population. Unstained cells were used to define background fluorescence, and gating for oxidized versus reduced C11-BODIPY populations was established using untreated controls.

### Immunofluorescence

Cells expressing the ALFA-tagged constructs were seeded on glass coverslips in 6-well plates. The next day, cells were washed with PBS and fixed in 4 % paraformaldehyde (Sigma) for 30 min, then quenched in PBS containing 0.1 M glycine for 10 min, and permeabilized and blocked for 15 min in PBS containing 10 % normal goat serum and 0.1 % Triton X-100. Coverslips were then incubated for one hour with FluoTag®-X2 anti-ALFA conjugated to Atto488 (diluted 1:500 in 3 % normal goat serum and 0.1 % Triton X-100). DNA was counterstained by incubating coverslips with DAPI (Thermo Fischer, 1:10000 in PBS) for 10 minutes, followed by two PBS washes for 10 minutes each. Coverslips were subsequently mounted onto microscope slides using DABCO (Sigma). Confocal fluorescence images were obtained using a Leica SP8 microscope with a HCX PL APO CS 63x 1.4 NA oil objective.

### Lipidomic analysis

Water-soluble metabolites were quantified by LC–MS. Briefly, 1 × 10□cells were harvested, washed with PBS, and snap-frozen in liquid nitrogen. Metabolites were extracted with 500 µL ice-cold methanol/water (80:20, v/v) supplemented with 0.01 µM lamivudine and 1 µM each of D□-glucose, D□-succinate, D□-glycine, and 1□N-glutamate (Sigma-Aldrich). After centrifugation, supernatants were evaporated and reconstituted in 150 µL of 5 mM NH□OAc in acetonitrile/water (50:50, v/v), and 20 µL aliquots were injected for analysis. Metabolites were separated on an XBridge Premier BEH Amide UPLC column (2.5 µm, 100 × 2.1 mm; Waters) using a DIONEX Ultimate 3000 UHPLC system. Solvent A was 5 mM NH□OAc in CH□CN/H□O (40:60, v/v) and solvent B was 5 mM NH□OAc in CH□CN/H□O (95:5, v/v). Separation was performed at a flow rate of 200 µL/min and at 45 °C using a linear gradient from 100% B to 10% B over 23 minutes, followed by 16 minutes in 10% B and a 2-minute return to 100% B. Columns were equilibrated with 100% B for 7 minutes before each injection. High-resolution mass spectrometry was performed on a Q Exactive mass spectrometer equipped with a HESI probe, operated in alternating positive and negative full MS mode (69–1000 m/z, 70K resolution; Thermo Fisher Scientific) using standard ESI parameters. Data were analyzed using TraceFinder™ v5.1, integrating peaks corresponding to the calculated monoisotopic masses of the metabolites (± H□, ± 3 mMU). All analyses were conducted in three independent biological replicates.

### Quantification and statistical analysis

Details of the statistical analyses for each experiment are provided in the corresponding figure legends, with *P* values indicated in the figures. Unless stated otherwise, data are shown as mean ± standard deviation (SD) from three independent biological replicates conducted on separate days. Statistical significance was assessed using one-way or two-way ANOVA followed by Dunnett’s or Tukey’s multiple-comparisons test (ns□=□not statistically significant, *p□<□0.05, * *p□<□0.01, * * *p□<□0.001, * * * *p□<□0.0001), performed in GraphPad Prism v10 (GraphPad Software). The outcomes of these analyses are reported in the figures. No statistical methods were used to predetermine sample size, and no data were excluded from the analyses. Experiments were not randomized, and investigators were not blinded to allocation during experimentation or outcome assessment.

### Reporting summary

Additional details on research design are available in the Nature Portfolio Reporting Summary linked to this article.

## Acknowledgements

J.P.F.A. acknowledges support from the Junior Group Leader program of Rudolf Virchow Center, University of Würzburg, and additional support from the Deutsche Forschungsgemeinschaft (DFG), FR 3746/3-1, FR 3746/5-1, FR 3746/6-1, CRC205 (INST 269/886-1), and TRR 387/1 (514894665); the EU-H2020 (ERC-Consolidator, DeciFERR); and the Deutsche Jose Carreras Leukämie Stiftung (DJCLS 01 R/2022). The São Paulo Research Foundation (Grants: #2023/04397-4, #2021/11597-4, #2022/03130-1). National Council for Scientific and Technological Development – CNPq (444300/2024-4).

## Conflict of Interest

The authors declare no conflict of interest.

## Data Availability Statement

All data and materials supporting the findings of this study are available within the main text, figures, and extended data figures. Uncropped blot images are presented in Supplementary Fig. 1. Additional information is available from the corresponding author upon reasonable request. Source data accompany this paper.

## Author Contributions

I.D.S. performed the majority of the experiments and drafted the manuscript. I.A.M.A., F.P.F., A.B.S.T., M.C.C.R., K.A.L., and A.F.S. contributed to methodology development. W.S. carried out the lipidomic analysis, and G.F. acquired the fluorescence microscopy images. J.P.F.A. and C.R.R.R. conceived the study, designed the experimental plan, supervised the work, and revised the manuscript. All authors contributed to data discussion, interpretation, and manuscript revision, and approved the final version.

## Notes

### Competing Interest Statement

The authors have declared no competing interest.

## References

1. Angeli, J.P.F., Shah, R., Pratt, D.A. & Conrad, M. Ferroptosis Inhibition: Mechanisms and Opportunities. Trends Pharmacol Sci 38, 489–498 (2017).

2. Dos Santos, A.F., Fazeli, G., Xavier da Silva, T.N. & Friedmann Angeli, J.P. Ferroptosis: mechanisms and implications for cancer development and therapy response. Trends Cell Biol (2023).

3. Dixon, S.J. & Olzmann, J.A. The cell biology of ferroptosis. Nat Rev Mol Cell Biol 25, 424–442 (2024).

4. Guo, Q. et al. The Role of Iron in Cancer Progression. Frontiers in Oncology 11(2021).

5. Buschhaus, J.M. et al. Effects of iron modulation on mesenchymal stem cell-induced drug resistance in estrogen receptor-positive breast cancer. Oncogene 41, 3705–3718 (2022).

6. Ru, Q. et al. Iron homeostasis and ferroptosis in human diseases: mechanisms and therapeutic prospects. Signal Transduct Target Ther 9, 271 (2024).

7. Galy, B., Conrad, M. & Muckenthaler, M. Mechanisms controlling cellular and systemic iron homeostasis. Nature reviews. Molecular cell biology 25, 133–155 (2024).

8. Yu, Y. et al. Hepatic transferrin plays a role in systemic iron homeostasis and liver ferroptosis. Blood 136, 726–739 (2020).

9. Brown, C.W. et al. Prominin2 Drives Ferroptosis Resistance by Stimulating Iron Export. Dev Cell 51, 575–586.e4 (2019).

10. Hassannia, B., Vandenabeele, P. & Vanden Berghe, T. Targeting Ferroptosis to Iron Out Cancer. Cancer cell 35, 830–849 (2019).

11. Chang, L.-C. et al.

12. Hassannia, B. et al. Nano-targeted induction of dual ferroptotic mechanisms eradicates high-risk neuroblastoma. Journal of Clinical Investigation 128, 3341–3355 (2018).

13. Zheng, C. et al. Donafenib and GSK[J4 Synergistically Induce Ferroptosis in Liver Cancer by Upregulating HMOX1 Expression. Advanced science 10, e2206798–n/a (2023).

14. Ren, T. et al. Zoledronic acid induces ferroptosis by reducing ubiquinone and promoting HMOX1 expression in osteosarcoma cells. Frontiers in pharmacology 13, 1071946–1071946 (2023).

15. Homan, R.A., Jadhav, A.M., Conway, L.P. & Parker, C.G. A Chemical Proteomic Map of Heme-Protein Interactions. J Am Chem Soc 144, 15013–15019 (2022).

16. Jeney, V.R. et al. Pro-oxidant and cytotoxic effects of circulating heme. Blood 100, 879–887 (2002).

17. Petrillo, S. et al. Heme accumulation in endothelial cells impairs angiogenesis by triggering paraptosis. Cell Death & Differentiation 25, 573–588 (2018).

18. Chiabrando, D., Vinchi, F., Fiorito, V., Mercurio, S. & Tolosano, E. Heme in pathophysiology: a matter of scavenging, metabolism and trafficking across cell membranes. Front Pharmacol 5, 61 (2014).

19. Immenschuh, S., Vijayan, V., Janciauskiene, S. & Gueler, F. Heme as a Target for Therapeutic Interventions. Front Pharmacol 8, 146 (2017).

20. Wu, J. et al. The non-canonical effects of heme oxygenase-1, a classical fighter against oxidative stress. Redox Biol 47, 102170 (2021).

21. Hsu, F.-F. et al. Signal peptide peptidase-mediated nuclear localization of heme oxygenase-1 promotes cancer cell proliferation and invasion independent of its enzymatic activity. Oncogene 34, 2360–2370 (2015).

22. Hori, R. et al. Gene Transfection of H25A Mutant Heme Oxygenase-1 Protects Cells against Hydroperoxide-induced Cytotoxicity. Journal of Biological Chemistry 277, 10712–10718 (2002).

23. Sun, J. et al. Heme regulates the dynamic exchange of Bach1 and NF-E2-related factors in the Maf transcription factor network. Proceedings of the National Academy of Sciences 101, 1461–1466 (2004).

24. Anandhan, A. et al. NRF2 controls iron homeostasis and ferroptosis through HERC2 and VAMP8. Science advances 9, eade9585–eade9585 (2023).

25. Tsuji, Y. et al. Coordinate Transcriptional and Translational Regulation of Ferritin in Response to Oxidative Stress. Molecular and Cellular Biology 20, 5818–5827 (2000).

26. Li, Y. et al. Inhibition of CISD2 promotes ferroptosis through ferritinophagy-mediated ferritin turnover and regulation of p62–Keap1–NRF2 pathway. Cellular & molecular biology letters 27, 81–81 (2022).

27. Goessling, L.S., Mascotti, D.P. & Thach, R.E. Involvement of Heme in the Degradation of Iron-regulatory Protein 2. Journal of Biological Chemistry 273, 12555–12557 (1998).

28. Jeong, J., Rouault, T.A. & Levine, R.L. Identification of a Heme-sensing Domain in Iron Regulatory Protein 2. Journal of Biological Chemistry 279, 45450–45454 (2004).

29. Consoli, V., Sorrenti, V., Gulisano, M., Spampinato, M. & Vanella, L. Navigating heme pathways: the breach of heme oxygenase and hemin in breast cancer. Molecular and Cellular Biochemistry (2024).

30. Fiorito, V., Chiabrando, D., Petrillo, S., Bertino, F. & Tolosano, E. The Multifaceted Role of Heme in Cancer. Front Oncol 9, 1540 (2019).

31. Fukuda, Y. et al. Upregulated heme biosynthesis, an exploitable vulnerability in MYCN-driven leukemogenesis. JCI Insight 2(2017).

32. Sohoni, S. et al. Elevated Heme Synthesis and Uptake Underpin Intensified Oxidative Metabolism and Tumorigenic Functions in Non-Small Cell Lung Cancer Cells. Cancer Res 79, 2511–2525 (2019).

33. Gozzelino, R., Jeney, V. & Soares, M.P. Mechanisms of cell protection by heme oxygenase-1. Annu Rev Pharmacol Toxicol 50, 323–54 (2010).

34. Zhu, X.G. et al. Functional Genomics In Vivo Reveal Metabolic Dependencies of Pancreatic Cancer Cells. Cell Metabolism 33, 211–221.e6 (2021).

35. Menon, A.V. et al. Excess heme upregulates heme oxygenase 1 and promotes cardiac ferroptosis in mice with sickle cell disease. Blood 139, 936–941 (2022).

36. Zille, M. et al. Hemin-Induced Death Models Hemorrhagic Stroke and Is a Variant of Classical Neuronal Ferroptosis. The Journal of Neuroscience 42, 2065–2079 (2022).

37. Hamamura, R.S. et al. Induction of heme oxygenase-1 by cobalt protoporphyrin enhances the antitumour effect of bortezomib in adult T-cell leukaemia cells. British Journal of Cancer 97, 1099–1105 (2007).

38. Yachie, A. et al. Oxidative stress causes enhanced endothelial cell injury in human heme oxygenase-1 deficiency. J Clin Invest 103, 129–35 (1999).

39. Alla, S.S.M. et al. A rare case of heme oxygenase deficiency: A case report and literature review. Clin Case Rep 12, e8986 (2024).

40. Yachie, A. Heme Oxygenase-1 Deficiency and Oxidative Stress: A Review of 9 Independent Human Cases and Animal Models. International Journal of Molecular Sciences 22, 1514 (2021).

41. Chau, A.S. et al. Heme oxygenase-1 deficiency presenting with interstitial lung disease and hemophagocytic flares. Pediatr Rheumatol Online J 18, 80 (2020).

42. He, J. et al. Reprogramming of iron metabolism confers ferroptosis resistance in ECM-detached cells. iScience 26, 106827 (2023).

43. Fang, X. et al. Loss of Cardiac Ferritin H Facilitates Cardiomyopathy via Slc7a11-Mediated Ferroptosis. Circulation Research 127, 486–501 (2020).

44. Deng, Y. et al.

45. Gurjao, C. et al. Discovery and Features of an Alkylating Signature in Colorectal Cancer. Cancer Discov 11, 2446–2455 (2021).

46. Bouvard, V. et al. Carcinogenicity of consumption of red and processed meat. Lancet Oncol 16, 1599–600 (2015).

47. Anderson, K.E. & Collins, S. Open-Label Study of Hemin for Acute Porphyria: Clinical Practice Implications. The American Journal of Medicine 119, 801.e1-801.e6 (2006).

