## Extended Data for "HMOX1 controls a heme–ferritin switch that protects cells from ferroptosis"

**Extended Data 1**

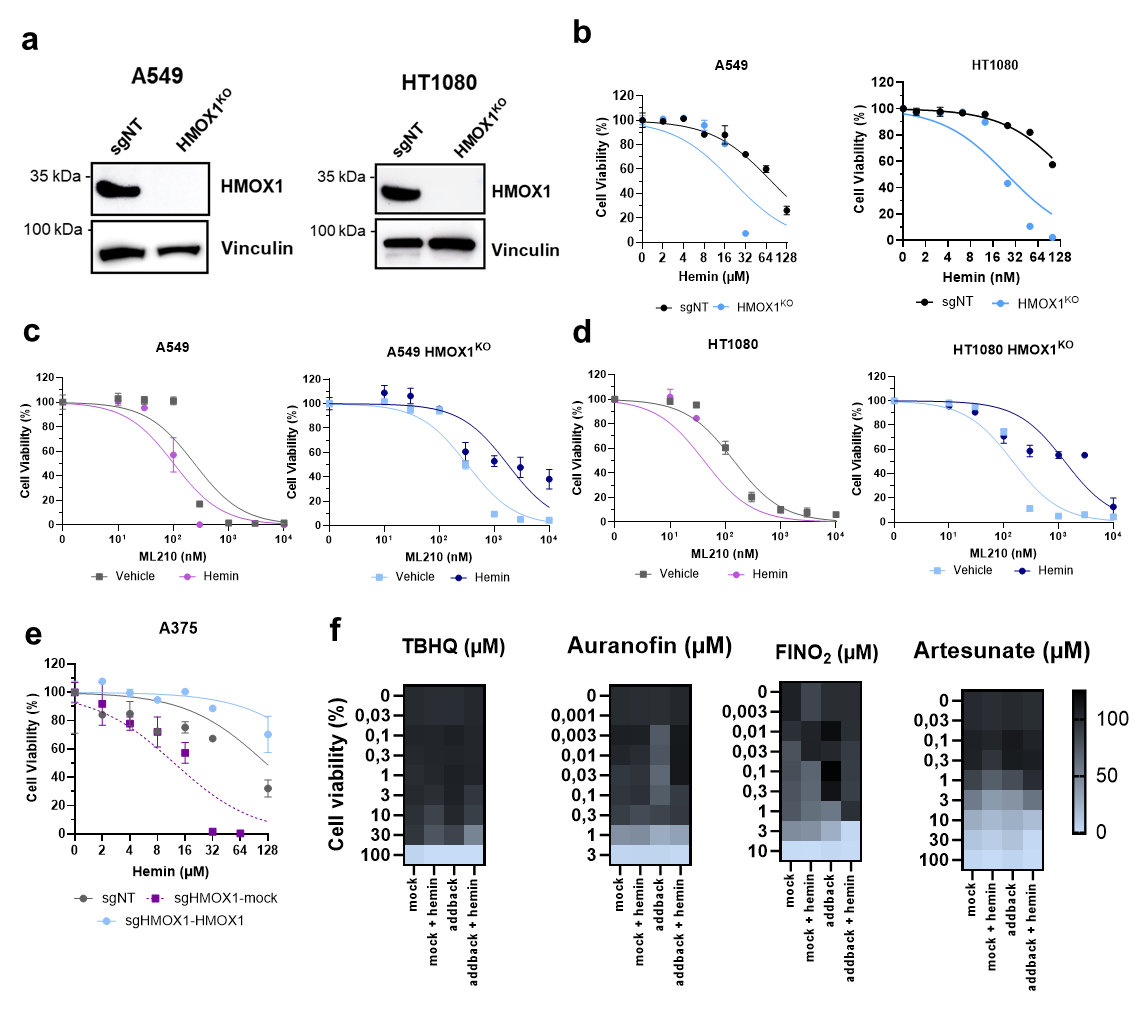

**Extended Data 1. a**, Immunoblot (IB) analysis of HMOX1, and vinculin in A549 and HT1080 sgNT and HMOX1^KO^ cells. **b**, Dose-dependent cytotoxicity in A549 and HT1080 sgNT and HMOX1KO cells treated with increased concentration of hemin (0 – 128 μM) for 72 h. **c**, Dose-dependent cytotoxicity in A549 sgNT and HMOX1KO cells pre-treated with hemin for 24 h (10 μM) followed by ML210 treatment for 72 h. **d**, Dose-dependent cytotoxicity in HT1080 sgNT and HMOX1KO cells pre-treated with hemin for 24 h (10 μM) followed by ML210 treatment for 72 h. **e**, Dose-dependent cytotoxicity in A375 sgNT and A375 HMOX1^KO^ cells reconstituted with either an empty vector control or a vector encoding HMOX1 cells treated with increased concentration of hemin (0 – 128 μM) for 72 h. **f**, Dose-dependent cytotoxicity in A375 HMOX1^KO^ cells reconstituted with either an empty vector control or a vector encoding HMOX1, after hemin pre-treatment (10 μM) followed by TBHQ, Auranofin, FINO2, and Artesunate (72 h). Cell viability was assessed using Alamar Blue. Data represent mean ± SD of triplicates from one representative experiment out of three performed.

**Extended Data 2**

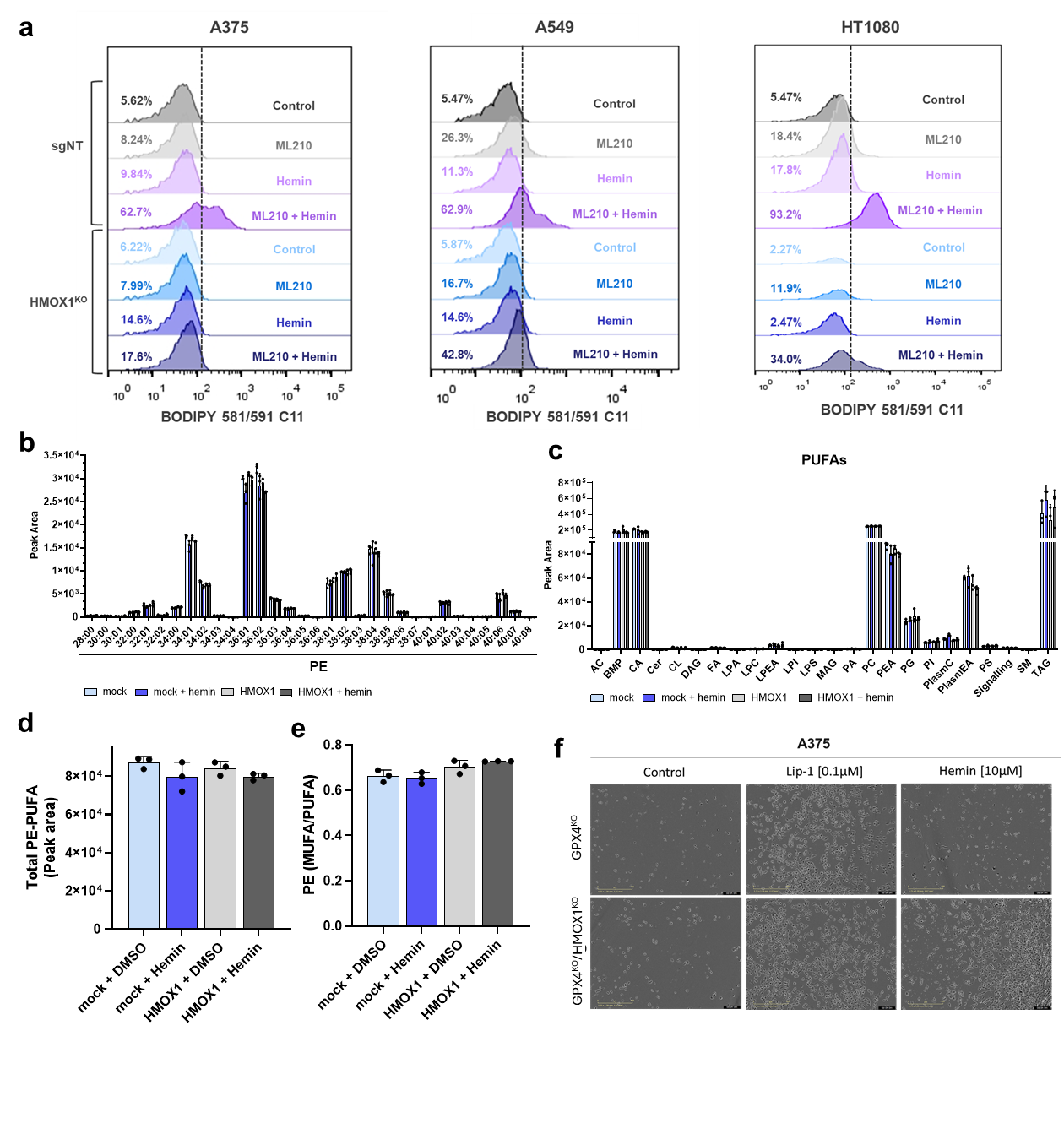

**Extended Data 2. a,** Lipid peroxidation analysis by C11-BODIPY 581/591 staining A375, A549, HT1080 sgNT and HMOX1KO cells pre-treated with hemin (10 μM, 24 h) and ML210 (300 nM, 4 h). Representative plots from one of two independent experiments are shown. **b**, Lipidomics analysis of A375 HMOX1^KO^ cells reconstituted with either an empty vector control or a vector encoding HMOX1 cell lines incubated for 24 h with hemin (10 μM). **b,d,** Presented are the total amount of PE containing PUFA, **c**, Fatty acid composition of PE species in A375 HMOX1^KO^ cells reconstituted with either an empty vector control or a vector encoding HMOX1 cell lines incubated for 24 h with hemin (10 μM). **e**, The ratio of mono- to polyunsaturated fatty acids (MUFA/PUFA) in PE species. **f**, A375 GPX4^KO^ and GPX4^KO^/HMOX1^KO^ cell lines were treated with hemin (10 μM, 24 h) or lip-1 (500 nM, 24 h) and cell growth was monitored using the IncuCyte imaging system.

**Extended Data 3**

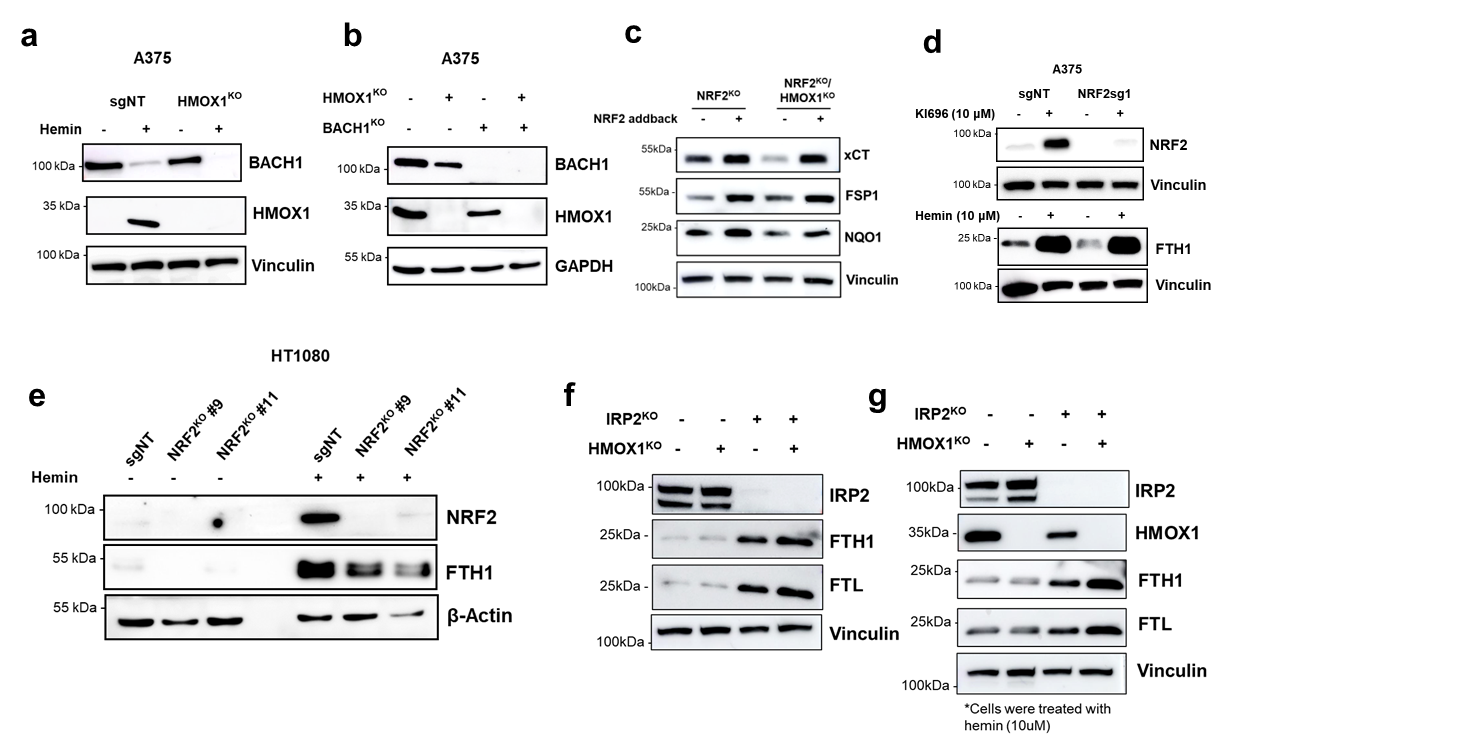

**Extended Data 3. a**, Immunoblot (IB) analysis of BACH1, HMOX1, and vinculin in A375 sgNT and HMOX1KO cells. **b**, Immunoblot (IB) analysis of BACH1, HMOX1, and GAPDH in A375 sgNT, HMOX1^KO^, BACH1^KO^ and HMOX1^KO^/BACH1^KO^ cells. **c**, Immunoblot (IB) analysis of xCT, FSP1, NQO1, and vinculin in A375 NRF2^KO^ and NRF2^KO^/HMOX1^KO^ cells reconstituted with NRF2. **d**, Immunoblot (IB) analysis of NRF2, FTH1, and vinculin in A375 sgNT and NRF2^KO^ treated with KI969 (10 μM, 24 h) and hemin (10 μM, 24 h). **e**, Immunoblot (IB) analysis of NRF2, FTH1, and β-Actin in HT1080 sgNT and NRF2^KO^ cell lines treated with hemin (10 μM, 24 h). **f**, Immunoblot (IB) analysis of IRP2, FTH1, FTL and vinculin in A375 sgNT, HMOX1^KO^, IRP2^KO^ and HMOX1^KO^/IRP2^KO^ cells. **g**, Immunoblot (IB) analysis of IRP2, HMOX1, FTH1, FTL and vinculin in A375 sgNT, HMOX1^KO^, IRP2^KO^ and HMOX1^KO^/IRP2^KO^ cells treated with (10 μM, 24 h).

**Extended Data 4**

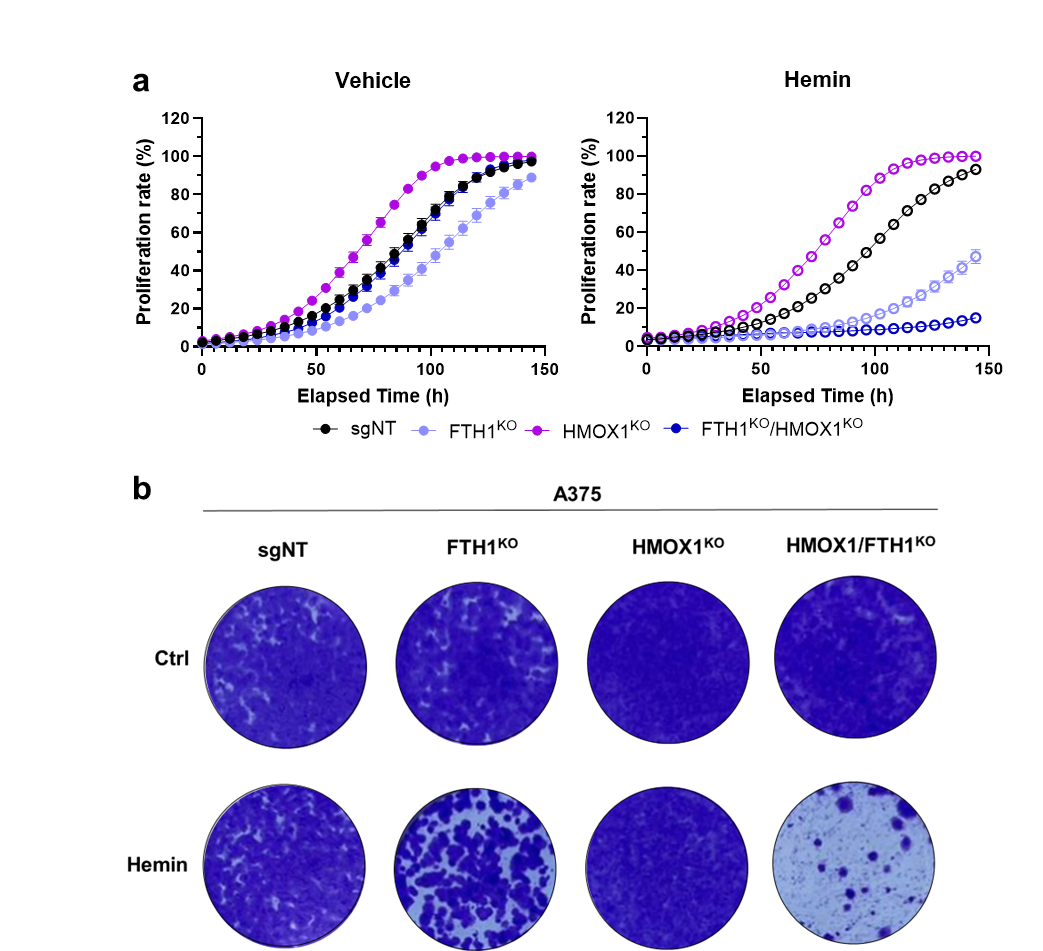

**Extended Data 4. a**, Cell growth analysis after hemin (10 μM) treatment in A375 sgNT, HMOX1KO, FTH1KO and HMOX1KO/FTH1KO cells. **b**, Crystal violet staining of A375 sgNT, HMOX1KO, FTH1KO and HMOX1KO/FTH1KO cells after 10 μM hemin treatment.
